# Cycling persister clones with elevated NR2F1-mediated cholesterol biosynthesis cause chemotherapy resistance

**DOI:** 10.64898/2026.06.06.730520

**Authors:** Hajime Sato, Taku Sato, Yohei Sasagawa, Ryoko Seki, Shicheng Zhang, Hiroshi Haeno, Daisuke Hishikawa, Mashito Sakai, Kenta Hata, Itoshi Nikaido, Yoshiyuki Mori, Tadahide Noguchi, Toshiaki Ohteki

**Affiliations:** Department of Biodefense Research, Medical Research Laboratory, Institute of Integrated Research, Institute of Science Tokyo (formerly Medical Research Institute, Tokyo Medical and Dental University (TMDU)), Tokyo 113-8510, Japan; Department of Dentistry, Oral and Maxillofacial Surgery, Jichi Medical University, Tochigi 329-0498, Japan; Department of Biochemistry and Molecular Biology, Nippon Medical School Graduate School of Medicine, Tokyo 113-8603, Japan; Department of Functional Genome Informatics, Biological Data Science, Medical Research Laboratory, Institute of Integrated Research, Institute of Science Tokyo, Tokyo 113-8510, Japan; Research Institute for Biomedical Science, Tokyo University of Science, Chiba 278-0022, Japan; Department of Oral and Maxillofacial Surgical Oncology, Graduate School of Medical and Dental Sciences, Institute of Science Tokyo, Tokyo 113-8510, Japan

**Author notes:** These authors contributed equally (Hajime Sato, Taku Sato).

## Abstract

While a subpopulation termed cycling persisters (CPs) that is characterized by sustained proliferation, even under chemotherapy drug exposure, contributes more directly to tumor relapse, the molecular basis for the emergence of CPs has been unclear. Here, we used the human tongue cancer organoid (TCO) library to continuously track the *in vitro* fate of individual cancer cell clones during and after chemotherapy exposure using time-lapse imaging. Among the heterogeneous clones, we identified CPs that formed larger clusters than the others, which barely grew and remained small (non-CPs). Using differences in cell cluster size as an indicator, thousands of CP and non-CP clones were directly sampled from 3D matrix organoid cultures and were analyzed. Notably, tumor-intrinsic interferon (IFN) signaling and hypoxic pathways were inactivated, whereas the NR2F1-mediated cholesterol biosynthesis pathway was distinctly activated in CPs compared to non-CPs. Indeed, inhibiting cholesterol biosynthesis with simvastatin significantly suppressed the appearance of CP clones, showing that elevated cholesterol biosynthesis is essential for the emergence of CPs. These findings suggest that clonal-level variations in the intensity of these signaling pathways determine the fate of individual tumor cells exposed to chemotherapeutic agents, which may provide insights into cancer relapse mechanisms and identify potential molecular targets of CPs.

## INTRODUCTION

While drug-tolerant persister (DTP) cells that arise after cytotoxic chemotherapy are in a diapause-like state with reduced metabolism and proliferation, they may not be entirely quiescent^1,2^. Given a subpopulation termed cycling persisters (CPs) that are characterized by sustained proliferation even under drug exposure and contribute more directly to tumor relapse^3^, understanding the molecular basis for the emergence and maintenance of CPs and elucidating their vulnerabilities, with a clear distinction between CPs and diapose-like DTPs, may provide important insights to prevent tumor recurrence and improve long-term treatment outcomes.

Unlike conventional cancer cell lines, patient-derived cancer organoids can maintain intratumor heterogeneity of the cancer cell clones in each patient’s tumor tissue^4–9^, which allows for the analysis of a diverse set of cancer cell clones. Previously, we generated a tongue cancer organoid (TCO) library of untreated TC patients with diverse cancer stages^10^. TCOs recapitulated the characteristics of the TC in each patient, such as their gene mutations and histological features^10^. Notably, each established organoid line had significantly different responsiveness to chemotherapy, and several lines with strong resistance were identified (chemo-resistant TCOs)^10^. Given that the exposure of chemo-resistant TCOs to cisplatin (cis-diaminedichloroplatinum (II), CDDP), a common agent used in TC chemotherapy, did not enrich CDDP-resistant clones, reversible drug tolerance, rather than drug resistance, is involved in the survival of TCOs during CDDP treatment^10^.

Here, we used the TCO library to determine whether CDDP treatment results in the emergence of CPs. We observed that the fate of persisters at the clonal level after CDDP treatment was heterogeneous, i.e., most died, some remained quiescent, and notably, some clones expanded. We established a method to distinguish CPs and non-CPs after CDDP exposure based on their morphological features under microscopy, and directly and separately sample those diverse clones from 3D matrix organoid cultures. Finally, we compared their gene expression profiles to characterize CPs.

Our study reveals a previously unappreciated mechanism of cancer relapse after chemotherapy and provides an opportunity to develop effective therapeutic strategies that target these mechanisms.

## RESULTS

### Clonal diversity in cell division is induced by chemotherapeutic agents

We used previously established TCO lines^10^ to observe the diversity of chemotherapy responsiveness in patients’ TC cells at the clonal level. In this study, we subjected four TCO lines we had already assessed for their responsiveness to chemotherapy: TCO8 and TCO21 were chemotherapy (chemo)-resistant lines, and TCO3 and TCO12 were relatively chemo-sensitive lines^10^. TCOs were dispersed into single cells and seeded into 3D Matrigel cultures in the absence (Fig. 1a, left) or presence of CDDP (Fig. 1a, right) and time-dependent changes in cluster formation from individual TC cells were observed under each condition. Cultured organoids were stained with calcein AM (for live cell staining) and propidium iodide (PI, for dead cell staining), after which 3D full-focus microscopic images were obtained (Fig. 1b). The area size of the individual surviving cell/cluster images was then quantified (Fig. 1c). In cultures without the addition of CDDP, on day 7 after the start of culture, the median area size of the TC cell cluster was increased by 15.7 times for TCO3, 9.0 times for TCO12, 14.3 times for TCO8, and 13.6 times for TCO21 compared to the median area size of each single TC cell at 24 h after seeding (Fig. 1b,c). This result confirmed that under these culture conditions, most chemo-resistant and chemo-sensitive TC clones underwent cell division and formed cell clusters.

**Figure 1.**
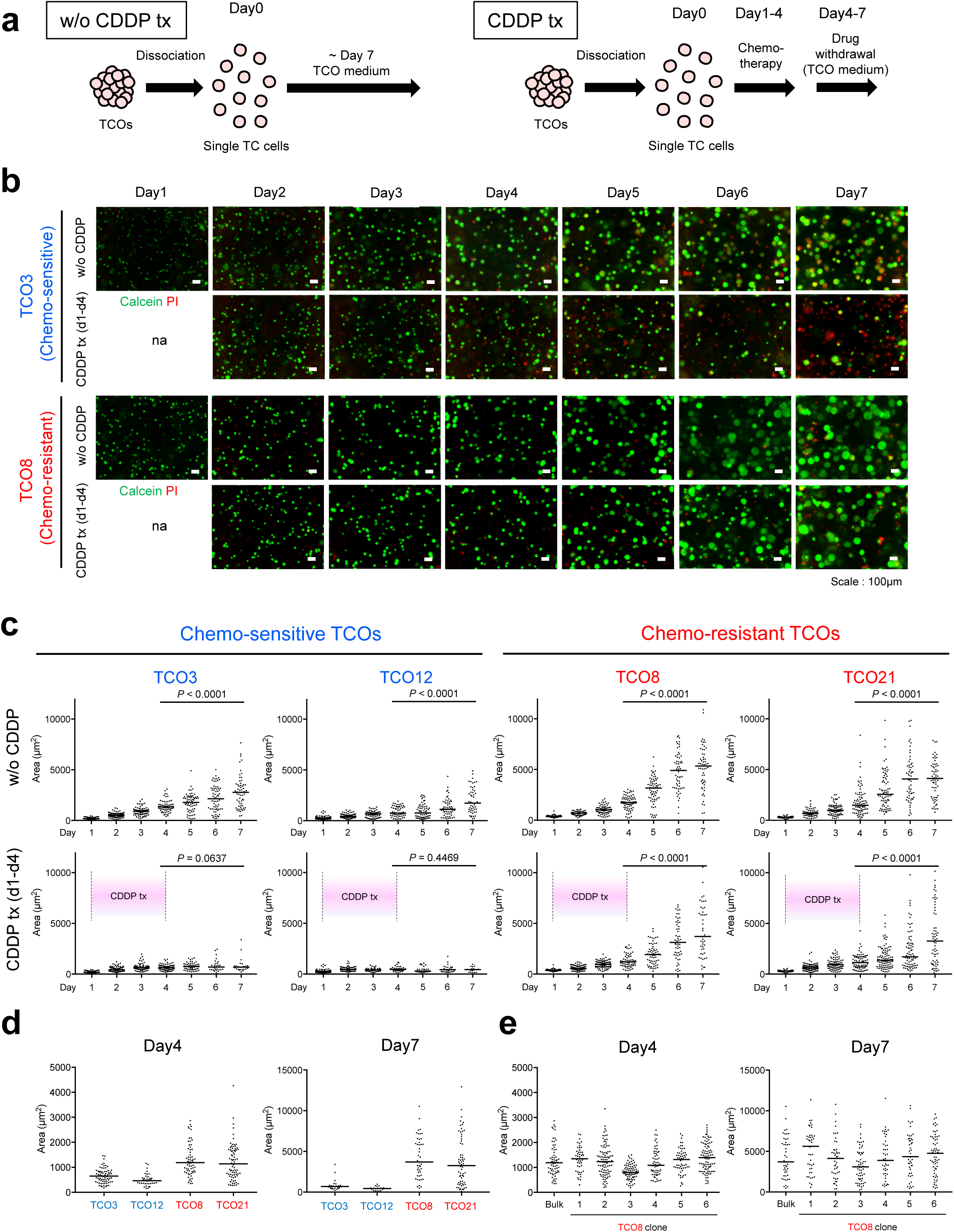
Clonal diversity in cell division is induced by chemotherapeutic agents. **a** Diagram of the experimental timeline. TCOs were dispersed into single cells, then plated and cultured without (left) or with (right) CDDP. CDDP treatment was performed from days 1 to 4, after which organoids were cultured in CDDP-free medium until day 7. **b** Representative images of TCO cultures. Chemo-sensitive TCO3 (top) and chemo-resistant TCO8 (bottom) were cultured with or without CDDP as in **a**. Fluorescence microscopic images were taken over time after staining with calcein AM (green, live cells) and PI (red, dead cells). Scale bars, 100 μm. **c** Quantification of area size in images of individual calcein AM-stained cell clusters. Statistical significance between data from days 4 and 7 was determined by Mann Whitney test. Horizontal bars indicate medians. Each experiment was performed two times, and similar results were obtained. n.s., not significant. **d** Comparison of the area size diversity between CDDP treated chemo-sensitive (TCO3, TCO12) and chemo-resistant TCO lines (TCO8, TCO21) TCO lines at days 4 and 7. **e** CDDP drug response in isogenic TC cells. Six single organoid clones were selected from bulk TCO8 and analyzed for cluster size diversity after CDDP exposure on days 4 and 7.

Next, we added CDDP to the TCO cultures from day 1 to day 4. The area size of individual clusters in images was continuously quantified until day 7 using the same method as above (Fig. 1a, right). Here, we used C_max_ as the CDDP concentration to set conditions close to those observed in patients. As a result, in the chemo-sensitive TCO lines (TCO3, TCO12), the size distribution of the viable cancer cell clusters on day 7 was comparable to that on day 4 (median size, TCO3 day 4: 646 µm^2^, TCO3 day 7: 688 µm^2^, TCO12 day 4: 465 µm^2^, TCO12 day7: 445 µm^2^). These results indicate that most of the viable TCO3 and TCO12 cancer cell clones entered a growth-arrested “cytostatic” state after CDDP exposure (hereafter referred to as “quiescent clones”) (Fig. 1b,c). We determined that the size distribution of quiescent clones at day 7 in TCO3 and TCO12 was 136-1060 µm^2^ (outlier clones were removed; see methods). On the other hand, when chemo-resistant TCO lines (TCO8, TCO21) were cultured and treated with CDDP, cancer cell clusters with various sizes had grown on day 7 (median size, TCO8 day 4: 1190 µm^2^, TCO8 day7: 3700 µm^2^, TCO21 day 4: 1137 µm^2^, TCO21 day 7: 3253 µm^2^) (Fig. 1b,c).

Similar to the case of the chemo-sensitive TCOs, the size distribution of all surviving clusters on day 4 was defined as the size of quiescent clones on day 7 in TCO8 and TCO21. Based on that definition, the sizes of quiescent clones in TCO8 and TCO21 were 488-2968 µm² and 121-2793 µm², respectively (outlier clones were removed; see Methods). Quiescent clones on day 7 in TCO8 and TCO21 accounted for an average of 37.4% (expt1: 39.1%, expt2: 35.7%) and 56.1% (expt1: 46.9%, expt2: 65.2%) of the total viable clones, respectively (Fig. 1c, Extended Data Fig. 1). On the other hand, in TCO8, an average of 62.6% (expt1: 60.9%, expt2: 64.3%) of clones analyzed on day 7 exceeded the size distribution observed on day 4, where such types of clones were defined as “expanded clones”. The size distribution of expanded clones in TCO8 was 3162-10536 µm^2^ (outlier clones were removed, Fig.1c). Similarly, in TCO21, an average of 44.0% (expt1: 53.1%, expt2: 34.8%) of clones analyzed were expanded clones (the size distribution of expanded clones: 3076-12931 µm^2^, Fig. 1c, Extended Data Fig. 1). Therefore, many cancer cell clones within chemo-resistant TCOs avoid entering dormancy and proliferate after CDDP removal. Such cell populations exhibit characteristics similar to cancer cells that proliferate during the “drug holiday” period following the end of chemotherapy in clinical settings^11^. Notably, the size diversity of cancer cell clusters was lost by CDDP treatment at day 4 in TCO3 and TCO12, whereas it was maintained substantially in the chemo-resistant TCOs. (Fig. 1d).

We next assessed whether TC cell survival during chemotherapy results from the diversity of genetic mutations that each clone possesses or is due to a non-genetic and transient response, such as drug tolerance. To confirm this, six organoid clones were randomly picked from chemo-resistant TCO8 using a microscope, dissociated into single cells, and cultured and expanded in an optimal medium with CDDP from day 1 to 4, where the cancer cells constituting each organoid clone have a uniform genetic background. On days 4 and 7, we evaluated the responsiveness of individual TC cells by measuring the size of the TCOs as an indicator. Notably, the size distribution of clusters formed by TC cells prepared from each organoid clone #1-#6 and the parental TCO8 line was comparable (Fig. 1e). Thus, the appearance of “expanded clones” is not due to the diversity of genetic mutations.

### Detection of CP clones during chemotherapy exposure

Cancer cells from chemo-resistant TCOs formed larger cell clusters during CDDP treatment than those from chemo-sensitive TCOs (Fig. 1d), which indicates that chemo-resistant TCOs evade the effects of chemotherapy and continue to proliferate. To confirm the cell division state of these clones, TC cell clusters were stained with calcein AM and Hoechst, after which the number of viable cells constituting individual clusters was counted under a microscope using the number of nuclei as an indicator.

At 24 hours after seeding, more than 95% of TC cell clones were single cells in all TCO lines tested (Fig. 2a,b). Furthermore, 72 hours after CDDP treatment (day 4 of culture), most TC cell clones derived from chemo-sensitive TCOs remained as single cells. In contrast, some clones derived from chemo-resistant TCOs formed cell clusters of 2–8 cells. (Fig. 2a,b), which indicated that CPs indeed emerged during chemotherapy treatment of chemo-resistant TCOs (Fig. 2c).

**Figure 2.**
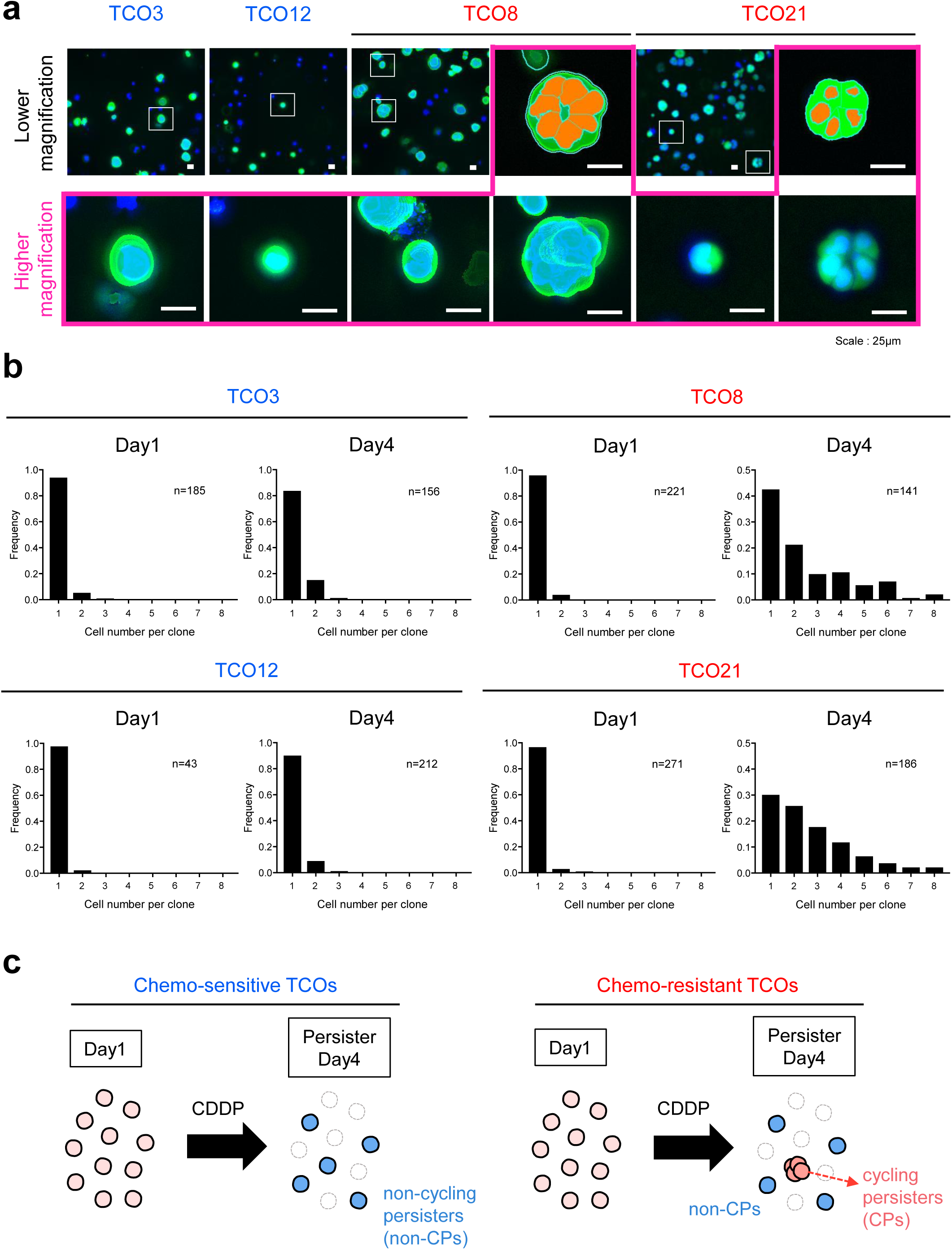
Detection of CPs under chemotherapy exposure. **a** Representative images of calcein AM/Hoechst 33342 staining at 72 hours (day 4) after exposure to CDDP in chemo-sensitive (TCO3, TCO12) or chemo-resistant (TCO8, TCO21) TCOs. Scale bars for high magnification images, 100 μm. Scale bars for low magnification images, 25 μm. **b** Quantification of cells constituting individual clusters based on Hoechst staining at day 1 (left) or day 4 (right) in the images shown in (a). The number n indicates the number of clusters analyzed. **c** Diagram showing persister cell status in TCOs after CDDP exposure. In chemo-sensitive TCOs (left), the majority of surviving cell clusters at day 4 were non-CPs. In contrast, chemo-resistant TCOs (right) contained not only non-CPs but also CPs that continued to divide after drug exposure and formed larger cell clusters.

Previous reports have shown that blocking the cell cycle by adding the CDK inhibitor abemaciclib to 3D cancer cell line organoid cultures has little effect on the frequency of persisters that survive in the presence of chemotherapy drugs, suggesting that persister survival is not solely due to cell cycle arrest^12^. Notably, the observation of CPs in this study supports the idea that cell cycle arrest is not the primary cause of persister survival.

### Clonal tracing after chemotherapy validates the potential of CPs

Because CPs occur stochastically, it is difficult to capture and analyze the process of CP formation or predict their future fate (e.g., whether they will expand or remain quiescent after drug removal). In this study, we focused on CPs that appeared during chemotherapy treatment and attempted to develop a method to identify them using a microscope.

Here, we performed time-lapse imaging to track the fate of individual single-cancer cells derived from TCOs after exposure to CDDP. We then focused on and analyzed the imaging characteristics of the “day 4” clones, which would become “expanded clones” on day 7. In this study, cancer cells derived from chemo-resistant TCO8 and TCO21 were seeded, and cancer cell clones (a total of 86 to 101 clones) that we randomly selected from multiple observation fields were monitored every 24 hours from day 4 to day 7 (Fig. 3a). The size of individual clones in the acquired images was then quantified at each time point (Fig. 3b), and changes over time were visualized (Fig. 3c).

**Figure 3.**
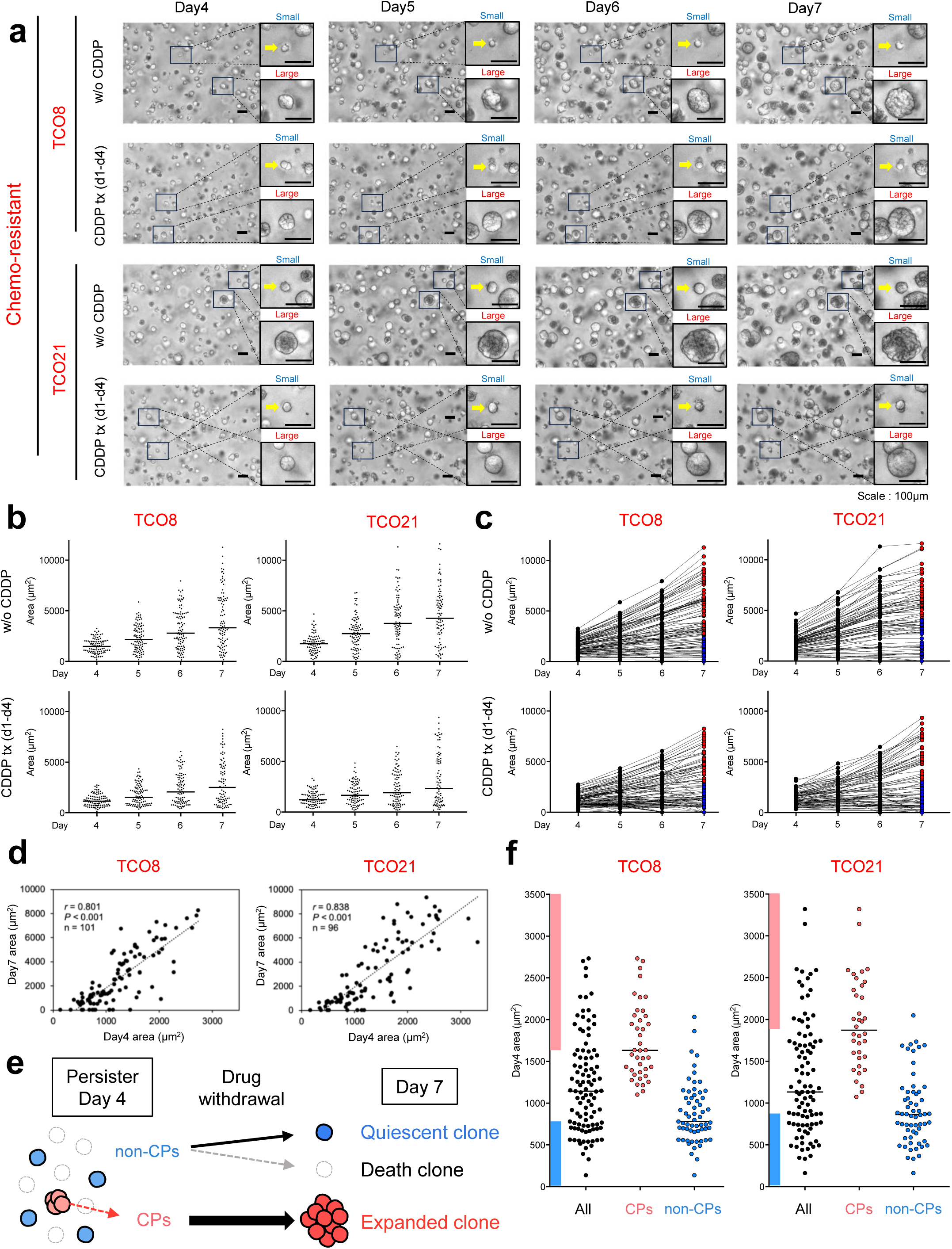
Clonal tracing after chemotherapy validates the potential of CPs. **a** Time-lapse series of bright-field microscopy images of chemo-resistant TCO lines (TCO8, TCO21) with or without CDDP treatment. In the CDDP-treated group, CDDP was added from day 1 to day 4, then further cultured in the absence of CDDP until day 7. Scale bars, 100 μm. **b** Quantification of the area size in images of individual clusters from day 4 to day 7 in individual tracked organoid clones (TCO8; n=101, TCO21; n=96). Each experiment was performed on biological replicates (n=2), and similar results were obtained in two experiments. **c** Size tracking of each organoid clone in (b) from day 4 to day 7. The red and blue symbols on day 7 indicate expanded clones and quiescent clones, respectively. **d** Correlation between area sizes of individual clusters on days 4 and 7. **e** Diagram showing the fate of persister cells after CDDP withdrawal. After drug withdrawal, CPs proliferated and formed larger cell clusters at day 7, called “expanded clones”. In contrast, non-CPs at day 4 either remained quiescent or died at day 7. **f** Size distribution of CP clusters (TCO8: n=40, TCO21: n=36) and non-CP clusters (TCO8: n=61, TCO21: n=60) identified by retrospective tracking of individual clones. CPs and non-CPs on day 4 were defined as clones that become expanded and quiescent clones, respectively, on day 7. The “All” group represents the size distribution of all individual clusters analyzed. Horizontal bars indicate medians in each group. Clusters in the size range between the CP and non-CP median had the potential to become both expanded and quiescent clones on day 7. Among CPs, red symbols represent clusters larger than the median CP, most of which became expanded clones on day 7, whereas among non-CPs, blue symbols represent clusters smaller than the median non-CP, most of which became quiescent clones on day 7.

As shown in Fig. 1, the size of the clusters at each time point was substantially diverse in both TCO8 and TCO21 cultured in the absence of CDDP (Fig. 3b, upper). Furthermore, tracking the changes in size of individual clones over time revealed that growth rates were highly variable across clones (Fig. 3c, upper). Similarly, when the chemotherapy drug CDDP was added, clones that formed larger cell clusters on day 4 became expanded clones on day 7 (the size distribution of “expanded clones”, TCO8: 3162-10536 µm^2^, TCO21: 3076-12931 µm^2^, Fig. 1c), but clones that formed relatively small clusters on day 4 either failed to grow or underwent cell death by day 7 (Fig. 3b,c, lower). Indeed, the cluster size on day 4 was positively correlated with the cluster size on day 7 both in TCO8 and in TCO21 (Fig. 3d). Therefore, in this culture system, the area values of the cancer cell clusters on day 4 during CDDP exposure can be used to predict which clones will become expanding or quiescent clones in the future (Fig. 3e).

The median areas of day 4 clusters that became “expanded clones” on day 7 (i.e., CPs) and day 4 clusters that became quiescent/death clones on day 7 (i.e., non-CPs) were 1634 µm^2^ and 789.5 µm^2^, respectively, in TCO8, and 1872 µm^2^ and 862.5 µm^2^, respectively, in TCO21 (Fig. 3f). Based on the size distribution of CPs and non-CPs in each chemo-resistant TCO calculated in Fig. 3f, we set a boundary value of the cluster size at which they could be accurately distinguished on day 4. Clusters in the size range between the CP and non-CP median could potentially become both expanded and quiescent/death clones on day 7. On the other hand, almost all cancer cell clusters larger than the median size of CPs grew into “expanded clones” by day 7, and nearly all the organoid clones smaller than the median size of non-CPs became quiescent/death clones by day 7 (Fig. 3f). Thus, this method identifies pure CPs in TCO8 and TCO21 lines under CDDP treatment.

### Establishment of a method to accurately harvest chemotherapy persister subclones

We next developed a method to directly harvest individual cancer cell clusters from 3D Matrigel cultures, enabling the comparison of gene expression profiles between CPs (which become expanded clones at day 7) and non-CPs (which become quiescent or dead clones at day 7). To this end, we used CELL HANDLER™ (YAMAHA) to harvest individual clusters (Fig. 4a). The success rates of collecting CPs and non-CPs (the number of harvested clusters/ the number of pickups) were 48.2%-98.8%, and the average success rate was 85.9%, confirming that the desired cancer cell clusters could be collected with high efficiency (Fig. 4b). With this sophisticated method, 72 hours after CDDP treatment (day 4 of culture), a total of 960 CPs (320 CPs/experiment x 3 experiments) were picked up from the cultures of TCO8 and TCO21. Furthermore, a total of 1440 non-CPs (480 CPs/experiment x 3 experiments) were picked up from the cultures of TCO8 and TCO21, and a total of 1380-1440 non-CPs (440-480 CPs/experiment x 3 experiments) were picked up from the cultures of TCO3 and TCO12 (Fig. 4c). The distribution frequency in cell cluster sizes, indicated by area (μm^2^), was similar in all three biological replicates (Fig. 4d).

**Figure 4.**
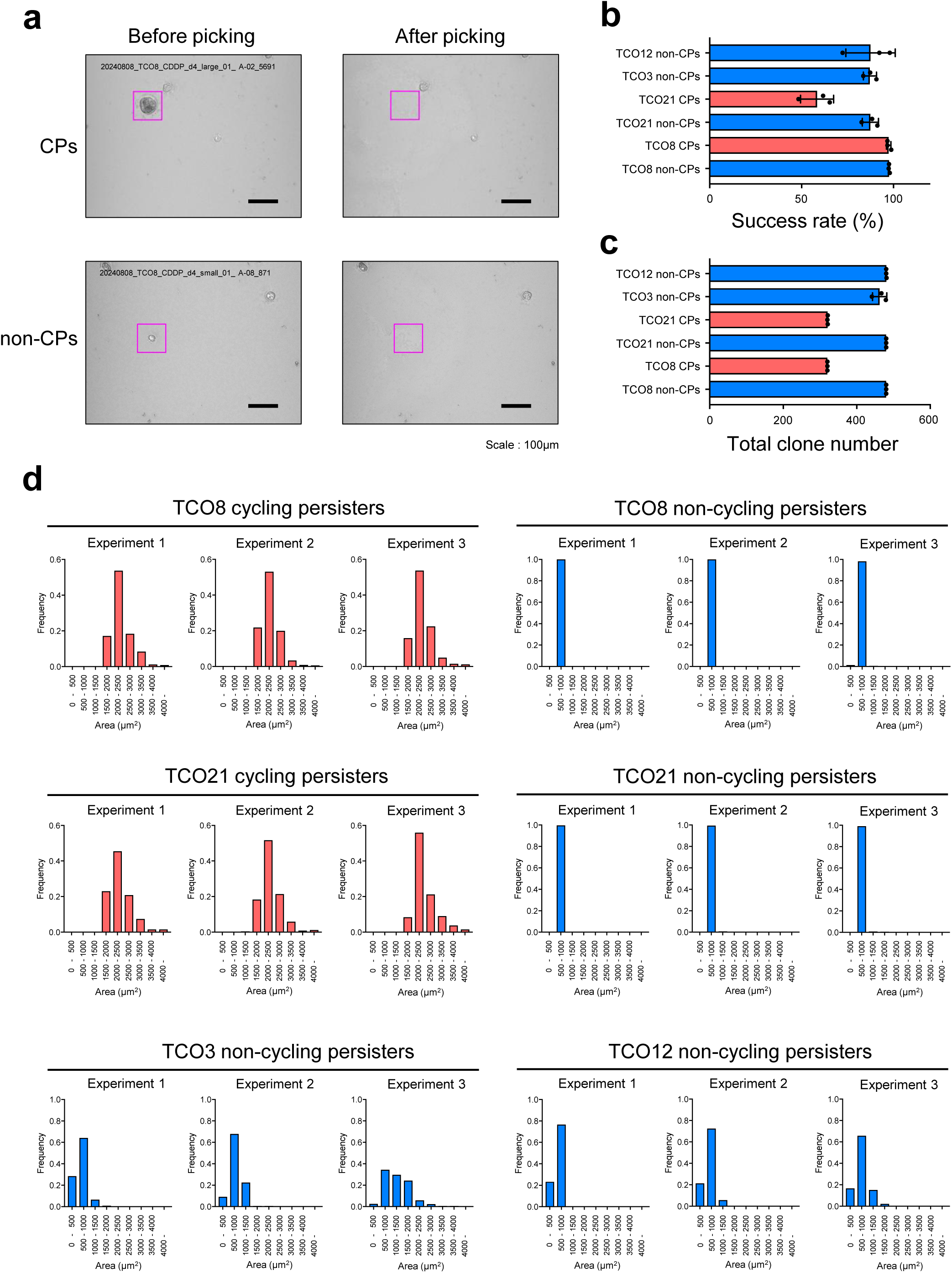
Establishment of a method to accurately harvest cycling persister subclones. **a** Representative brightfield images of organoids before (left) and after (right) harvesting using a CELL HANDLER™ system. TCO3, TCO12, TCO8 and TCO21 were treated with CDDP from day 1 to day 4 (72 hours), after which CPs and non-CPs were harvested based on the size of the clusters. Scale bars, 100 μm. **b** Success rates (%) of harvesting CPs and non-CPs, calculated by dividing the number of harvested clusters by the number picked up using the CELL HANDLER^TM^. Data are reported as means ± s.d. of three experiments as biological replicates. **c** Total numbers of CP and non-CP clones harvested. Data are reported as means ± s.d. of three experiments as biological replicates. **d** Size distribution of harvested CPs and non-CPs. The x-axis represents the distribution of cluster sizes, and the y-axis shows the frequency of picked clusters of each size. For TCO8 and TCO21, clusters larger than the median CPs (TCO8; 1632.5 µm^2^, TCO21; 1872 µm^2^), most of which became expanded clones on day 7, were harvested for gene expression analysis as “CPs”, whereas clusters smaller than the median non-CPs (TCO8; 780 µm^2^, TCO21; 862.5 µm^2^), most of which became quiescent/death clones on day 7, were harvested for gene expression analysis as “non-CPs”. For TCO3 and TCO12, clusters were harvested randomly because most of those clusters were non-CPs, as shown Fig. 1c.

### Gene expression profiles in CPs and in non-CPs during chemotherapy treatment

In this study, we aimed to clarify the gene expression advantages of CPs compared to non-CPs, thereby elucidating the mechanism by which CPs are formed and maintained. Thus, RNA-seq analysis was performed on CPs derived from TCO8 and TCO21 and non-CPs derived from TCO8, TCO21, TCO3, and TCO12, all harvested as shown in Fig. 4. Hierarchical clustering and PCA revealed that inter-line differences in gene expression among individual TCO lines were greater than those between CPs and non-CPs (Fig. 5a,b). At first, these gene expression profiles were compared to assess the differential expression of candidate genes responsible for chemotherapeutic drug resistance. It is well established that CDDP resistance occurs when the expression of several transporters involved in CDDP uptake and efflux is altered^12,13^. However, the expression of most of those transporter genes was low. Even if they were expressed, little change was seen between CPs and non-CPs within the same TCO line (Extended Data Fig. 2). DNA repair enzymes for platinum drug adducts, including *ERCC1, XRCC1, XRCC2, XRCC3,* and *XPA*, are also known to affect the sensitivity to platinum-based chemotherapy^13^. However, no differences in the expression of those genes were found between CPs and non-CPs (Extended Data Fig. 3). Therefore, it is unlikely that any of those factors are involved in the emergence of CPs in CDDP-treated TCOs.

**Figure 5.**
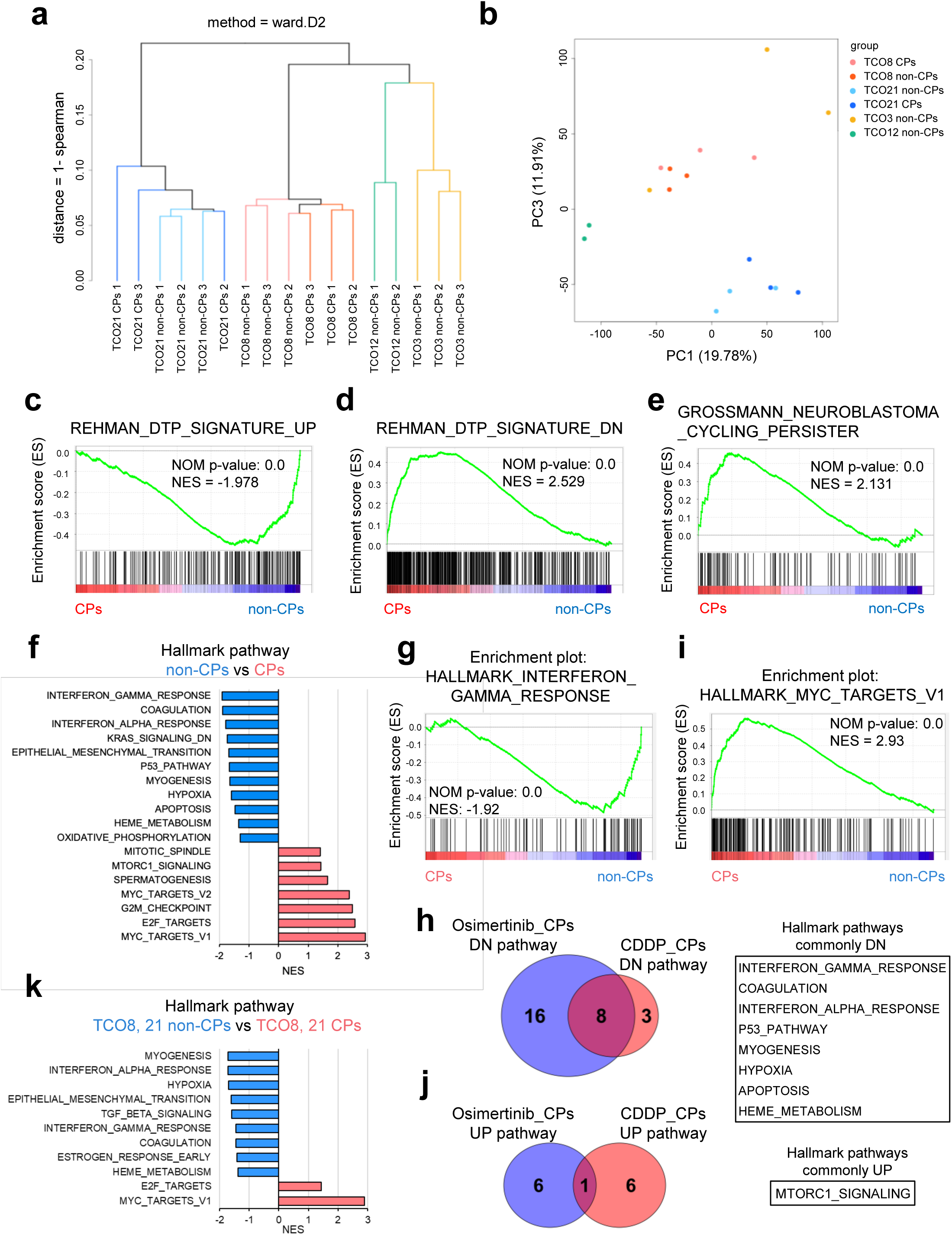
Gene expression profiles in CPs and in non-CPs during chemotherapy treatment. **a** Dendrogram obtained from the hierarchical clustering of average Spearman correlations showing transcriptome differences between CPs from TCO8 and TCO21, and non-CPs from TCO8, TCO21, TCO3, and TCO12. **b** PCA plot showing gene expression variation among samples. **c** GSEA plot showing enrichment of the gene set upregulated in diapause-like DTPs (REHMAN_DTP_SIGNATURE_UP; ref. 2) in non-CPs. **d** GSEA plot showing enrichment of the gene set downregulated in diapause-like DTPs (REHMAN_DTP_SIGNATURE_DN; ref. 2) in CPs. **e** GSEA plot showing enrichment of the gene set upregulated in CP cells from neuroblastoma patients (GROSSMANN_NEUROBLASTOMA_CYCLING_PERSISTER; ref. 3). **f** Significantly different hallmark pathways between CPs (from TCO8 and TCO21) and non-CPs (from TCO8, TCO21, TCO3, and TCO12) were detected by GSEA. Significantly different pathways were determined when the NOM p-value was less than 0.05. The graph was ordered by normalized enrichment score. **g** GSEA plots showing the most enriched gene sets in non-CPs of panel **f**. **h** Venn diagram showing the overlap of significantly activated hallmark pathways between osimertinib-induced CPs (ref. 14) and CDDP-induced CPs (this study). Pathways were identified as significantly activated in each group compared to their respective controls. The shared hallmark pathways are listed on the right. **i** GSEA plots showing the most enriched gene sets in CPs of panel **f**. **j** Venn diagram showing the overlap of significantly inactivated hallmark pathways between osimertinib-induced CPs (ref. 14) and CDDP-induced CPs (this study). Pathways were identified as significantly inactivated in each group compared to their respective controls. The shared hallmark pathways are listed on the right. **k** Significantly different hallmark pathways between CPs (from TCO8 and TCO21) and non-CPs (from TCO8 and TCO21) were detected by GSEA. Significantly different pathways were determined when the NOM p-value was less than 0.05. The graph was ordered by the normalized enrichment score. NES: normalized enrichment score., NOM p-value: nominal p-value.

Next, Gene Set Enrichment Analysis (GSEA) was performed based on gene expression profiles between chemotherapy-induced CPs, previously reported diapause-like DTPs from experimental tumor models^1,2^, and CPs identified in tumor tissues from chemotherapy-treated neuroblastoma patients^3^. That analysis showed that gene sets that were upregulated in diapause-like DTPs relative to untreated controls were significantly enriched in non-CP population (Fig. 5c). Conversely, gene sets that were downregulated in diapause-like DTP tumors were preferentially enriched in CPs identified in our study (Fig. 5d), indicating that non-CPs, but not CPs, closely resemble the diapause-like DTPs. Moreover, gene sets characteristic of CPs identified in neuroblastoma were significantly upregulated in CPs compared to non-CPs (Fig. 5e). These results suggest that CPs identified in our study more closely resemble the proliferative CPs that cause neuroblastoma relapse^3^ than the cytostatic diapause-like DTPs^1,2^.

To identify pathways characteristic of CPs and non-CPs, we further analyzed their gene expression profiles and found that CPs exhibited marked downregulation of several biological pathways, including “INTERFERON_GAMMA_RESPONSE”, “COAGULATION”, “INTERFERON_ALPHA_RESPONSE”, “P53_PATHWAY”, “MYOGENESIS”, “HYPOXIA”, “APOPTOSIS”, and “HEME_METABOLISM”, compared to non-CPs (Fig. 5f,g). Notably, most of those pathways were also significantly downregulated in CPs identified in neuroblastoma^3^. Consistently, Oren and colleagues reported similar suppression of those pathways in CPs of an EGFR-mutated NSCLC cell line after osimertinib treatment^14^ (Fig. 5h). These observations suggest that the inactivation of these pathways may represent a shared requirement for establishing the CP state, independently of the cancer type or drug class. As expected, in CPs identified in this study, pathways primarily associated with cell proliferation, such as “MYC_TARGET_V1,” “E2F_TARGET,” and “G2M_CHECKPOINT” were upregulated (Fig. 5f,i). On the other hand, activation of the antioxidant response or fatty acid oxidation pathways, which were previously reported to be necessary for the survival and maintenance of CPs of EGFR-mutated NSCLC cells^14^, was not observed in CPs identified in this study. In this context, “MTORC1_SIGNALING” was the only pathway commonly upregulated in CPs in this study and in those described by Oren and colleagues^14^ (Fig. 5f,j). Therefore, the activation pathways involved in driving the transition into the CP state appear to vary depending on the cell type and treatment context. In line with the pan-line analysis, similar changes in biological pathways were also observed when CPs and non-CPs were compared within the same chemo-resistant TCO line (Fig. 5k).

In conclusion, our results suggest that the balance between activated and suppressed signaling pathways at the clonal level may determine the fate of individual tumor cells in response to chemotherapeutic agents.

### Persistent NR2F1-mediated transcriptional priming of the cholesterol biosynthesis pathway is required for the formation of CPs

In our previous study, we found that chemo-resistant TCOs exhibit constitutive activation of the cholesterol biosynthesis pathway compared to chemo-sensitive counterparts, suggesting a strong link between this pathway and chemo-resistance^10^. Importantly, this characteristic was highly enriched in the CPs following CDDP treatment. This was also the case within the same chemo-resistant TCO lines (Fig. 6a). These results suggest that clonal differences in the basal activation level of the cholesterol biosynthesis pathway may determine the fate of each cancer cell upon chemotherapy exposure, namely, whether the cell enters a cytostatic state or survives while retaining proliferative capacity. Supporting this notion, combined treatment with CDDP and simvastatin, an inhibitor of HMGCR, the rate-limiting enzyme in the cholesterol biosynthesis pathway, significantly reduced the number of CPs and their subsequent expanded clones compared to CDDP treatment alone (Fig. 6b,c).

**Figure 6.**
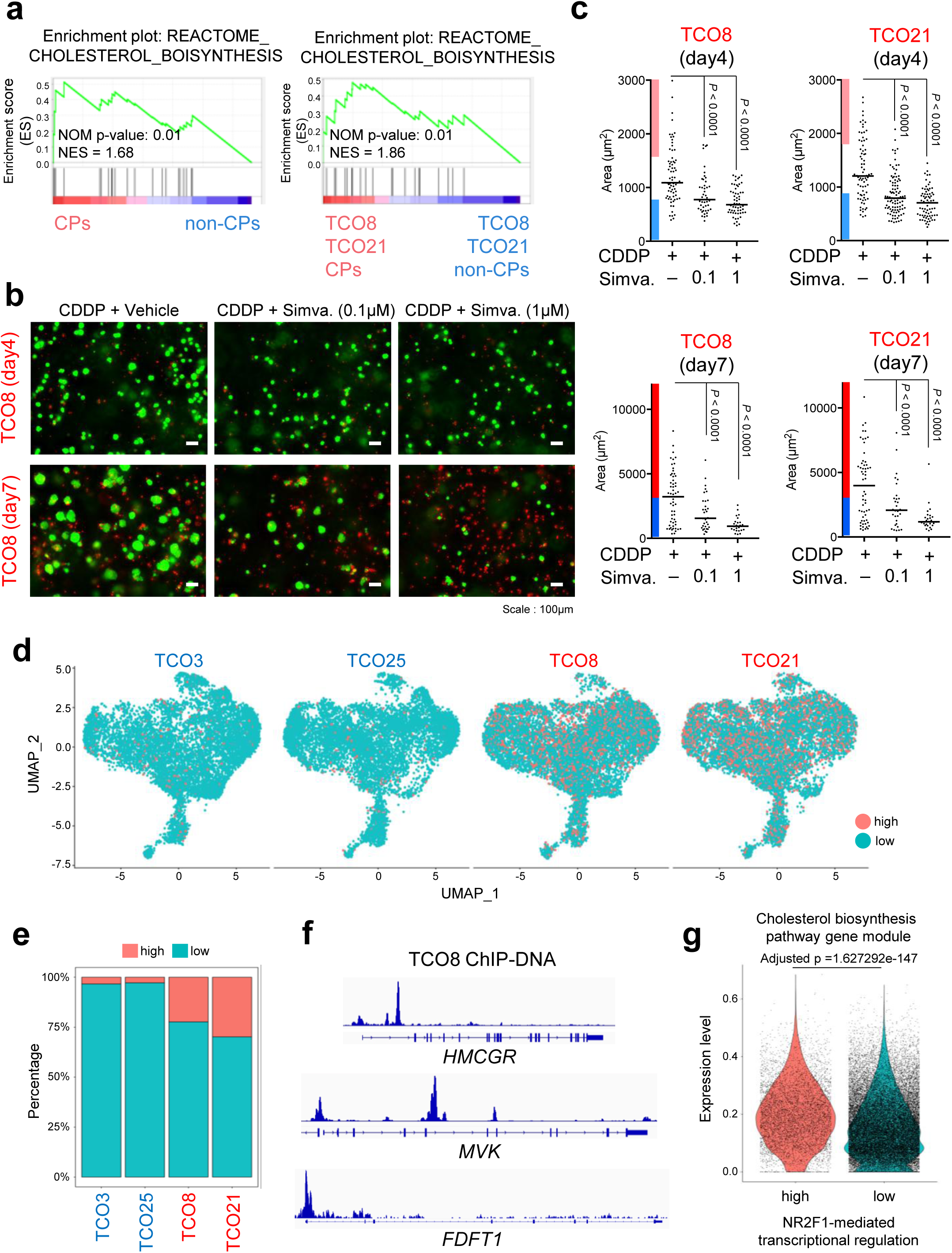
CPs are potentially induced by the activation of the cholesterol biosynthesis pathway via NR2F1. **a** GSEA plots of cholesterol biosynthesis between CPs from TCO8, TCO21, and non-CPs from TCO8, TCO21 or TCO8, TCO21, TCO3, and TCO12. NES: normalized enrichment score. NOM p-value: nominal p-value. **b** Representative images of TCO8 at day 4 or day 7 of culture in the presence of 10 μM CDDP, with or without Simvastatin (0.1 or 1 μM). Fluorescence microscopic images were taken after staining with calcein AM (green, live cells) and PI (red, dead cells). Scale bars, 100 μm. **c** Quantification of area size in images of individual calcein AM-stained cell clusters in TCO8 and TCO21 at the endpoint (day 7). Statistical significance was determined by one way ANOVA with multiple comparison tests. Each experiment was performed on biological replicates (n=2), and similar results were obtained in two experiments. **d** UMAP plot based on scMultiome data, showing cells with both high *NR2F1* gene expression (top 25%) and high chromatin accessibility at its binding motif (orange) versus cells with low expression/accessibility (blue). **e** Stacked bar plot quantifying the proportion of NR2F1-high (orange) and NR2F1-low (blue) cells in each TCO line (TCO3, 12, 8, 21), as defined in (**d**). **f** Representative IGV tracks showing ChIP-seq signals of NR2F1 at genomic loci of cholesterol biosynthesis pathway genes (*HMGC*, *MVK*, and *FDFT1*). Enrichment of NR2F1 at these loci confirms its direct association with regulatory regions of these genes. **g** Violin plots showing expression levels of cholesterol biosynthesis pathway-related genes in NR2F1-high and NR2F1-low cells defined in (**d**) and (**e**). Statistical analysis was performed using “MAST” method with Bonferroni correction in the “FindMarkers” function in “Seurat” package.

To investigate how such cholesterol-dependent clones emerge under steady-state conditions, we reanalyzed our previously acquired scMultiome dataset^10^. Specifically, we focused on NR2F1, the only transcription factor whose chromatin accessibility was elevated in chemo-resistant TCOs in our earlier analyses^10^. In Fig. 6d, the cells highlighted in orange on the UMAP plot exhibited both high expression of the NR2F1 gene (top 25%) and high chromatin accessibility at its binding motif, as assessed by chromVAR deviation scores (Fig. 6d,e). Notably, the NR2F1-transcription-activated cells were markedly enriched in the chemo-resistant TCOs, indicating a potential role of NR2F1 in the establishment or maintenance of the chemo-resistant state.

To assess whether NR2F1 constitutively binds to loci related to the cholesterol biosynthesis pathway, we performed ChIP-seq in untreated chemo-resistant TCO8 and TCO21, followed by comprehensive quality control and peak analysis. Global ChIP-seq metrics, including the number of called peaks, total mapped reads, and FRiP scores, indicated a high signal-to-noise enrichment (Extended Data Fig. 4a). Peak annotations revealed that NR2F1 predominantly binds to intronic, distal intergenic, and promoter regions (Extended Data Fig. 4b). Representative IGV views confirmed robust NR2F1 binding at typical peak regions, with clear enrichment over input controls (Supplementary Fig. 4c). ChIP-seq signals were highly reproducible across biological replicates, as shown by strong Pearson correlations (Extended Data Fig. 4d). In addition, de novo motif analysis confirmed enrichment of the known NR2F1 binding motif at peak centers (Extended Data Fig. 4e), supporting the specificity of NR2F1-DNA interactions. These results collectively demonstrate that the ChIP-seq data for NR2F1 are of high quality, with high specificity and reproducibility.

Importantly, we observed prominent NR2F1-binding peaks at multiple loci of cholesterol biosynthesis pathway genes, including *HMGCR*, *MVK*, and *FDFT1*, suggesting direct transcriptional regulation (Fig. 6f). Consistently, scMultiome RNA-seq analysis revealed that NR2F1-transcription-activated cells exhibited a significantly elevated expression of genes involved in the cholesterol biosynthesis pathway (Fig. 6g, Extended Data Fig. 5). These findings suggest that persistent NR2F1-mediated transcriptional priming of the cholesterol biosynthesis pathway underlies the survival and maintenance of chemo-resistant clones.

## DISCUSSION

Several studies have shown that CPs emerge occasionally when cancer cells are exposed to molecular targeted drugs. Oren et al. established an advanced method that can track the fate of individual cancer cells using a DNA barcoding system and can simultaneously measure their proliferation and transcriptional status. Using this system, they measured the behavior of EGFR-mutated NSCLC cell lines after treatment with the EGFR inhibitor osimeritinib and found that 13% of persisters were highly proliferative and could form large colonies. Notably, those CPs exhibited an enhanced antioxidant stress response or fatty acid oxidation pathways^14^. On the other hand, the exposure of BRAFV600E mutant melanoma cell lines to BRAF inhibitors *in vitro* demonstrated that 5% of persisters were CPs exhibiting sporadic ERK signaling activation^15^. These reports indicate that treatment of cancer cell lines with molecular targeted drugs induces stochastic changes in gene expression and signaling activity at the clonal level, leading to the emergence of CPs. In contrast, cytotoxic chemotherapeutic drugs have been thought to induce DTPs with a diapause-like phenotype, characterized by a quiescent or slow-cycling state and a suppressed metabolic activity^1,2^. However, recent single-nucleus RNA sequencing analyses of tumor tissues from chemotherapy-treated patients have revealed the presence of CPs even after chemotherapy, although their abundance varies between individuals^3^. Notably, a higher prevalence of CPs correlates with significantly shorter patient survival, underscoring their potential role in driving tumor relapse^3^. While these observations from clinical samples provide critical insights, they offer limited information regarding the molecular mechanisms by which CPs survive chemotherapy. Therefore, the development of *in vitro* models that reliably recapitulate the chemotherapy-induced emergence of CPs is essential to uncover the biological underpinnings of CPs and to identify potential therapeutic vulnerabilities. In this regard, using comparative analysis of TCOs that reproduce patient-specific cancer characteristics and cancer cell heterogeneity, we discovered that chemo-resistant TCOs exhibited an enrichment of CPs following chemotherapy treatment, whereas chemo-sensitive TCOs did not. The CPs grew into large organoids during and after drug removal. In contrast, non-CPs barely grew and remained small. As transcriptomic features, the tumor cell-intrinsic interferon signaling and hypoxia-related pathways were downregulated in CPs compared to non-CPs. Notably, these features were similar to those reported in CPs from osimertinib-treated EGFR-mutant NSCLC cells^14^ and chemotherapy-treated neuroblastoma tumors (Fig. 5e), suggesting a common transcriptomic program across CPs. Accordingly, tumor-intrinsic IFN signaling is well known to inhibit the growth of bone-disseminated dormant prostate tumors^16^. Thus, a release from growth inhibition via intrinsic IFN signaling may stochastically induce CPs (Fig. 5f,g,k). Another finding was that hypoxic signaling was characteristically suppressed in CPs (Fig. 5f,k). In this connection, we have previously shown that chromatin accessibility of HIF1A, which is critical for hypoxia-inducible gene expression, is constitutively higher in tumor cells of chemo-sensitive TCO3 and TCO12 than in chemo-resistant TCO8 and TCO21, while HIF1A mRNA expression levels are comparable. Thus, the HIF1A transcriptional program is activated explicitly in chemo-sensitive TC cells^10^. In the case of TC, the constitutive activation of the cholesterol biosynthesis pathway may play a critical role in maintaining the proliferative capacity of CPs even under chemotherapy. However, this feature has not been observed in CPs induced in cancer cell line models, suggesting that it represents a novel characteristic. This finding highlights the limitations of cell line-based analyses, which fail to capture the intratumoral heterogeneity of cancer clones. In contrast, the use of patient-derived tumor organoids, which can recapitulate the diverse clonal architecture of patient tumors, enables the identification of previously unrecognized properties and vulnerabilities of CPs.

Sustained activation of the cholesterol biosynthesis pathway via the transcription factor NR2F1 may represent a potential mechanism underlying this process. Although unrelated to the proliferation mechanism of CP, NR2F1, along with several other transcription factors, is also associated with transcriptional programs governing cholesterol metabolism in the colorectal neoplasia of *BRAF^V600E^*-mutated mice^17^. While NR2F1 has been implicated in inducing dormancy in disseminated tumor cells (DTCs) via epigenetic reprogramming^18,19^, our findings indicate that NR2F1 does not promote dormancy. Supporting this notion, NR2F1 was also identified as one of the defining genes in neuroblastoma CPs^3^, strongly suggesting that it plays a role distinct from that of previously reported dormancy-associated genes.

DTPs emerge in response to cytotoxic chemotherapy but it remains to be determined whether these cells pre-exist prior to treatment. For example, Ohta et al. used *in vitro* and *in vivo* imaging systems in mouse xenograft models of patient-derived colon cancer organoids to reveal that persister cancer cells exist as a subpopulation of colon cancer stem cells even before chemotherapy treatment^20^. Using a DNA barcoding system in PDX models or in 3D cultured organoids derived from colon cancer patients, other studies have shown that specific cancer cell clones were not enriched within the cells that survived chemotherapy treatment, suggesting that, at least in this experimental system, certain chemotherapy-resistant persister cells do not exist before drug treatment^1,2^. Furthermore, a mathematical model analysis showed that persistence can occur equally in all cells^1,2^. Consistent with the latter model, our system also demonstrated that when multiple subclonal lines derived from the same parental organoid were exposed to cytotoxic chemotherapy, both CPs and non-CPs emerged at comparable frequencies (Fig. 1e). This finding supports the idea that the appearance of persisters is induced by drug exposure rather than being pre-existing. Notably, the observation that the frequency of CP emergence varies significantly between different organoid lines strongly suggests the presence of intrinsic, line-specific factors that influence the probability of CP generation. In this study, we identified constitutive activation of the cholesterol biosynthesis pathway mediated by NR2F1 as one such heritable predisposition that facilitates the emergence of CPs.

Our study utilized a CELL HANDLER^TM^ that can capture the intra-tumor diversity of cancer cell clones including CPs and non-CPs at specific time points. This method is particularly useful for analyzing various patient-derived cancer organoid models at a clonal level. Although CPs can be considered the immediate and primary drivers of tumor recurrence following chemotherapy^14^, the possibility that non-CPs or slow-CPs may also serve as a source of relapse over longer time scales cannot be excluded. These non-CP populations may remain quiescent or minimally proliferative for extended periods before being reactivated under favorable conditions. Therefore, further characterization of non-CPs remains for future study, including elucidation of their survival duration, the environmental or cellular cues that trigger their re-entry into the cell cycle, and their potential contribution to tumor regrowth.

## METHODS

### Organoid culture

For organoid cultures, single cells were embedded in a 25μL drop of Matrigel (Corning) dome/well and seeded into 48-well plates. After the Matrigel solidified, 250 μL of TCO culture medium^10^ (AdDF+++, 1× B27 (Invitrogen), 1.25 mM N-acetylcysteine (Sigma-Aldrich), 100 ng/mL Human R-spondin1 (Peprotech), 100 ng/mL noggin (Miltenyi), 50 ng/mL EGF (Peprotech), 10 µM Y-27632 (Nacalai) and 50 μg/mL Primocin) was added to each well. Cells were cultured at 37 °C with 5% CO2, and the medium was changed every 3-4 days. For passaging, organoids were collected and incubated in 0.05% Trypsin/EDTA at 37 °C for 12 min. Following pipetting, the dissociated single cells were washed with 10% FCS/PBS, after which they were centrifugated at 1,500rpm for 5 min and the supernatant was removed. Cell pellets were washed with AdDF+++ and centrifuged again at 1,500 rpm for 5 min. After removing the supernatant and counting cells, single cells were suspended in Matrigel and seeded into 48-well plates. Organoids were passaged every 7-10 days depending on their proliferation rate.

For cryopreservation, dissociated cells were resuspended in CryoScarless DMSO-Free (Bio Verde) and were then stored at −80°C.

For CDDP treatment of TCOs, CDDP was added from days 1 to 4 after which they were cultured in CDDP-free medium until day 7.

### Organoid imaging

For fluorescence imaging of cancer cell clusters, 2μM Calcein-AM (live cell staining, Nacalai, Cat#19177-14) and 1μg/mL propidium iodide (dead cell staining, PI, Sigma-Aldrich Cat#25535-16-4) or 1 μ g/mL Hoechst 3332 (cell nuclei staining, Nacalai, Cat#19172-51) were added to the culture medium and incubated for 30 minutes at room temperature in the dark. Cell cluster images were taken daily from days 1 to 7 using a BZ-X810 microscope (Keyence). For analysis of image data, organoid areas in the images were measured using a BZ-X800 Analyzer with BZ-H4C software (Keyence), and each TCO line was analyzed in two independent experiments.

For time-lapse imaging, cell cluster images were taken over time within the same field of view from day 4 to day 7 using a BZ-X810 microscope (Keyence) with a stage top incubator (TOKAI HIT). TIn this system, cultures were maintained at 37°C with 5% CO2 during imaging. For analysis of image data, organoid areas in the images were measured using a BZ-X800 Analyzer (Keyence), and each TCO line was analyzed in two independent experiments.

To define “expanded clones” and “quiescent/death clones (Q/D clones)”, we identified outliers in the area data on day 4 using the interquartile range (IQR). Clones with sizes larger than the maximum value (excluding outliers) on day 7 were defined as “expanded clones”, while those with smaller sizes were described as “Q/D clones.”

### Organoid picking

After dissociation into single cells, cancer cells were suspended in Matrigel and sparsely seeded in 24-well plates (200-500 cells/50 μL Matrigel/well). They were treated with CDDP from day 1 to day 4, after which both CPs and non-CPs were harvested into 0.2 mL Hi-8 tubes (Takara) containing 10 μL culture medium using a CELL HANDLER™ (YAMAHA). A total of 320 CPs and 440-480 non-CPs per sample were collected in three independent experiments. The precision tip chip was replaced with a new one after every four collections, and images were captured before and after collection to confirm the accurate targeting of the clusters. Immediately after collection, trizol solution (a 3:1 mixture of TRIzol-LS and RNA-free water) was added to each PCR tube in a volume equivalent to three times the total volume of the collected organoids with culture medium, which included organoids, Matrigel, and any residual collected solution. Each collected organoid was estimated to increase by approximately 0.36 μL per organoid collection, and this additional volume was taken into account when calculating the required trizol solution. The solution in each PCR tube was then mixed for 1 minute at 4°C, and then centrifuged at 2,000 rpm using a ThermoMixerC with SmartBlock PCR 96. After centrifugation, the samples were stored at −80°C.

### RNA-seq library preparation

Organoids dissolved in trizol in multiple 0.2 ml tubes were thawed at room temperature and then combined into one or two 1.5 ml DNA LoBind Tubes for each sample. The solution was centrifuged by spin-down to remove the pellet containing the Matrigel, after which the supernatant was collected in one 5 ml DNA LoBind Tube and measured for volume. An equal volume of 99.5% EtOH was added to the obtained supernatant in the 5 ml tube and was mixed well by inverting mixing. The mixed solution was used to obtain purified total RNA using ta Direct-zol RNA Microprep Kit (Zyomo Research). The purified total RNA solution was checked for RNA quality by High Sensitivity RNA ScreenTape Analysis (Agilent).

Next, to prepare a sequence library DNA for next-generation sequencing (NGS) from the purified, minute amount of RNA, a GenNext® Shin-RamDA-seq® Single Cell Stranded Kit (TOYOBO, Cat#RML-101T) was used.

Following the addition of dilution buffer (consisting of Lysis Buffer, Lysis Enhancer, and RNase Inhibitor) to the purified, minute quantity of RNA, a denaturation step was performed by incubating the mixture at 70°C for 1.5 minutes. To eliminate genomic DNA, a genome removal premix (consisting of RT-RamDA® Buffer, gDNA Remover, rRNA Remover, and nuclease-free water) was added, followed by incubation at 30°C for 5 minutes. Subsequently, an RT-RamDA premix (consisting of RT-RamDA® Buffer, RT-RamDA® Enzyme Mix, 1st NSR Primer Mix for humans (NSR Primer Set for humans, Cat# NSR-101), and nuclease-free water) was added to the reaction mixture. Each mixture was then incubated under the following conditions: 25°C for 10 min, 30°C for 10 min, 37°C for 30 min, 50°C for 5 min, and 98°C for 5 min to allow for cDNA synthesis and amplification. Subsequently, the second strand synthesis was carried out by adding a second strand synthesis premix (consisting of 2nd strand synthesis Buffer, 2nd strand synthesis Enzyme, and 2nd NSR Primer Mix for human (NSR Primer Set for humans, Cat# NSR-101)) and incubating it at 16°C for 60 min and then at 80°C for 15 min.

To purify double-stranded cDNA, a purification procedure was carried out. The cDNA was mixed with 1/4 diluted Agencourt AMPure XP beads (Beckman Coulter) and subjected to magnetic bead-based purification using 80% ethanol. After washing and drying, the purified cDNA was eluted with 10 mM Tris-HCl (pH 8.0).

Subsequently, the purified double-stranded cDNA was subjected to fragmentation, end repair, and A-tailing by adding a premix containing Fragmentase, End Repair and A-tailing Buffer, and End Repair and A-tailing Enzyme. The reaction was performed by incubation at 30°C for 30 minutes followed by 65°C for 5 minutes.

To perform adapter ligation, IDT for Illumina-TruSeq UD Indexes v2 (96 Indexes, 96 Samples, Cat# 20040870) was diluted to a final concentration of 50 nM in 10 mM Tris-HCl (pH 8.0) while maintaining the temperature below 10°C to prevent degradation. The diluted adapter solution and ligation solution were added to each sample after fragmentation, end repair, and A-tailing. Unique index adapters were used for each sample to enable multiplexing. The ligation reaction was carried out by incubating the mixture at 2,000 rpm for 2 minutes at 4°C on a ThermoMixer, followed by centrifugation at 10,000 g for 1 minute at 4°C. Subsequent thermal cycling was performed at 25°C for 10 min, 30°C for 10 min, 37°C for 30 min, 50°C for 5 min, and 98°C for 5 min. After ligation, the products were purified using 1× Agencourt AMPure XP beads. After purification, PCR amplification was performed using a PCR solution premix (Library Amplification Master Mix、Library Amplification Primer Mix) to enrich the ligated library. The PCR conditions were as follows: 94°C for 30 sec, followed by 19-20 cycles of 98°C for 10 sec, 60°C for 10 sec, and 68°C for 15 sec.

To purify the sequence library, magnetic bead-based purification was performed using 1× Agencourt AMPure XP beads and 80% ethanol. After washing and drying the beads, the purified library was eluted with 10 mM Tris-HCl (pH 8.0). The purified sequence library DNA was then collected from the supernatant using a magnetic stand and then stored at −80°C.

### RNA-seq and data analysis

The libraries were submitted to an Illumina NovaSeqX plus (Illumina), and the paired-end (2×150 bp) sequencing was performed by CyberomiX Inc. The reads were aligned to the reference human genome (hg38) with Hisat2 (v2.2.1, RRID: SCR_015530) software. The mapped reads were assigned to genes using FeatureCounts from the Bioconductor (RRID: SCR_006442) package Rsubread (v2.0.6, RRID: SCR_016945). A hierarchical clustering dendrogram of samples was generated using hclust with average linkage (UPGMA) on a distance matrix defined as 1 − Spearman’s rank correlation (ρ) between gene expression profiles. Analyses were performed in R 4.3.2 (RRID:SCR_001905). PCA was performed using R software (v4.3.2, RRID:SCR_001905) with the ‘prcomp’ function. The PCA plot was generated using ggplot2 (v3.4.4, RRID: SCR_014601).

### GSE analysis

GSEA^21,22^ was performed using GSEA v4.3.3 software (Broad Institute, RRID: SCR_003199), and the Hallmark gene set of MSigDB (h.all.v7.2.symbols.gmt) was used for analysis. The number of permutations was set to 1000. Lowly expressed genes were excluded in the GSEA datasets.

### Single-cell multiome preprocessing and integration

Raw NovaSeq reads were aligned using Cell Ranger ARC v2.0.2 against the 2020-A reference (Ensembl 98 + GENCODE v32). Gene-expression matrices entered Seurat (R) and were filtered to cells containing 1 000–50 000 UMIs, ≥ 500 genes and < 40 % mitochondrial transcripts^23^. Corresponding ATAC fragments were processed in Signac^24^: peaks were called with MACS2^25^ on standard chromosomes (GenomeInfoDb^26^) and intersected with the ENCODE (RRID: SCR_006793) blacklist^27^ via GenomicRanges::reduce^28^. Peaks < 20 bp or > 100 kb were discarded; cells were retained when fragments numbered 1 000–100 000, enriched peaks > 500, nucleosome signal < 4 and TSS enrichment > 3.

For cross-donor scRNA integration, matrices were normalized and variance-stabilized with SCTransform^29^, which includes PCA (100 PCs) and the SCT anchor pipeline (SelectIntegrationFeatures, PrepSCTIntegration and FindIntegrationAnchors) to generate a harmonized expression manifold. scATAC integration paralleled the Signac vignette: informative peaks (FindTopFeatures) underwent TF-IDF scaling and SVD to yield uncorrected LSI embeddings, which were anchored with FindIntegrationAnchors (reduction = “rlsi”) and corrected through IntegrateEmbeddings. Motif annotation employed AddMotifs with JASPAR 2024^30^ and per-cell activities were computed with RunChromVAR^31^. Integrated scRNA data were re-projected (100 PCs), a Euclidean k-nearest-neighbor graph was built (FindNeighbors) and clusters were identified at resolution 0.3 (FindClusters); global structure was visualised with UMAP (RunUMAP).

### Gene-set scoring in single cells

To characterize transcriptional programs, curated GMT files from MSigDB^21,22^ were read with read.gmt() (clusterProfiler, RRID: SCR_016884). Signature activity per cell was quantified with AddModuleScore_UCell from UCell^32^, enabling rapid, rank-based enrichment assessment independent of library size.

### ChIP-seq Library Preparation and Sequencing for NR2F1

Cells (1–2 × 10⁶) were crosslinked with 2mM disuccinimidyl glutarate (DSG, ProteoChem) at room temperature for 30 minutes, followed by additional crosslinking with 1% formaldehyde (Thermo Fisher) for 10 minutes at room temperature. The reaction was then quenched with 120 mM glycine, and the cells were washed with cold PBS. Chromatin was extracted and sonicated using a M220 Ultrasonicator (Covaris) to achieve DNA fragments of 200–500 bp. Immunoprecipitation was conducted overnight at 4°C using an antibody against transcription factor NR2F1 (PERSEUS PROTEOMICS, #H8132) and Protein A magnetic beads (Thermo Fisher). After reversal of crosslinking and purification, ChIP and input DNA were used to prepare sequencing libraries using the NEBNext Ultra II DNA Library Prep Kit (New England Biolabs) according to the manufacturer’s protocol. Libraries were sequenced using the AVITI™ system (Element Biosciences) to generate 50-bp single-end reads.

### NR2F1 ChIP-seq peak calling and motif analysis

Four NR2F1 ChIP replicates plus inputs were pooled and processed with MACS3 (--nomodel, 147 bp) to produce narrowPeak and pile-up tracks. Signal tracks for log₂-fold-enrichment and –log₁₀ p values were derived with macs3 bdgcmp, sorted, bgzipped and indexed; differential peaks were called with macs3 bdgpeakcall (cut-off = 2, length ≥ 200 bp, gap ≤ 100 bp). Per-peak mean signals were appended with BEDTools map^33^. Replicate concordance was confirmed by deepTools (RRID: SCR_016366, multiBamSummary bins, 10 kb; plotCorrelation)^34^. FASTA windows (±250 bp) were extracted with bedtools getfasta. Peaks were further annotated with HOMER (RRID: SCR_010881, annotatePeaks.pl; findMotifsGenome.pl)^35^, and the JASPAR 2024 CORE vertebrate PWMs provided the background motif library.

All motif-flagged peak sets were imported into R as GRanges. Genomic context was assigned with ChIPseeker^36^. Gene-level maxima were ranked for functional analysis: Hallmark MSigDB GSEA and GO-BP enrichment were executed with clusterProfiler^37^.

### Experimental Design

No randomization, blinding, or formal power analysis was performed in this study. Experimental groups were assigned according to predefined treatment conditions. Clones were selected randomly for imaging. All outcome measurements were based on objective or quantitative assays, minimizing the risk of bias.

### Protocol and Code Information

All experimental procedures are described in detail in the Methods section, providing full step-by-step protocols within the manuscript. Custom code used in this study can be shared upon reasonable request to the corresponding author.

### Statistical Analysis

Statistical analyses were performed using R software (v4.3.2, RRID:SCR_001905) and GraphPad Prism v.7 (RRID:SCR_002798).

Spearman’s rank correlation between paired, ordinal-scale organoid sizes at different time points was calculated using the R function cor.test(), assuming a monotonic relationship and no significant outliers.

Differences between two independent groups were assessed using a two-tailed Mann-Whitney U test. Data were at least ordinal and assumed to come from populations with identical shapes. Statistical significance was defined as a p-value less than 0.05.

## RESOURCE AVAILABILITY

### Lead contact

Futher information and requests for resources should be directed to and will be fulfilled by the lead contact, Dr. Toshiaki Ohteki.

### Data availability

The RNA-seq datasets have been deposited in the Gene Expression Omnibus (RRID: SCR_005012) with the accession number GSExxxxxx.

## ACKNOWLEDGEMENTS

We thank H. Kamioka for secretarial support and Mr. Saburo Ito for his valuable support in optimizing the cell collection conditions using the Cell Handler system. This work was supported by JSPS Grant-in-Aid for Challenging Research (Pioneering) under Grant Number JP21K18259 (T.S.), Takeda Science Foundation (T.S.), the Uehara Memorial Foundation (T.S.), Project Mirai Cancer Research Grants (T.S.), a collaborative research with YAMAHA MOTOR CO., LTD. (Y.S.), JSPS KAKENHI: grant number 22H04925 (PAGS) (T.S.), Nanken-Kyoten, Science Tokyo (Y.M., T.N.), and the Medical Research Center Initiative for High Depth Omics, TMDU (T.O.).

## AUTHOR CONTRIBUTIONS

H.S. and T.S. contributed equally to this work. T.S. conceived the study. H.S., T.S., Y.S., R.S., D.H., M.S., and K.H. performed experiments and analyzed data. H.S., T.S., Y.S., H.H., M.S., and T.O. wrote the manuscript. T.S., S.Z., and H.H. performed bioinformatic analyses. Y.S., M.S., H.H., I.N., Y.M., and T.N. provided tongue tissue samples, advice, and discussion. T.O. supervised the overall project.

