## Extended Data 1-5 for "Cycling persister clones with elevated NR2F1-mediated cholesterol biosynthesis cause chemotherapy resistance"

#### Extended Data Fig. 1. Sato *et al.*

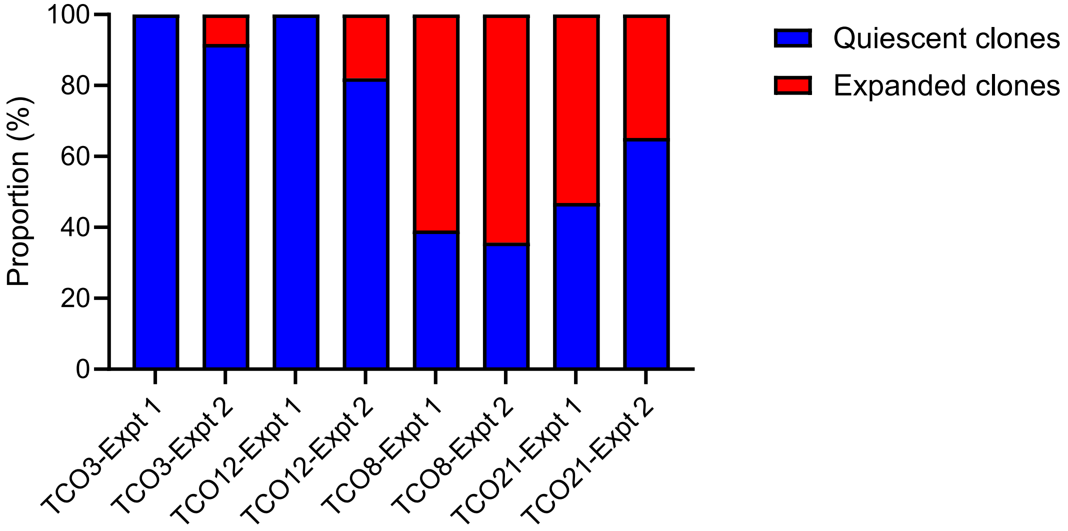


**Extended Data Fig. 1. Expanded clones emerged in chemo-resistant TCOs after chemotherapy treatment (related to Fig. 1c).** TCO lines (TCO3, TCO12, TCO8, and TCO21) were treated with CDDP from day 1 to day 4 and then cultured without the drug until day 7. Stacked bar plots show the proportions of quiescent clones (blue) and expanded clones (red) in each TCO line at day 7. Each TCO line was analyzed in two independent experiments, shown as Expt1 and Expt2.

**Extended Data Fig. 2. Sato *et al.***

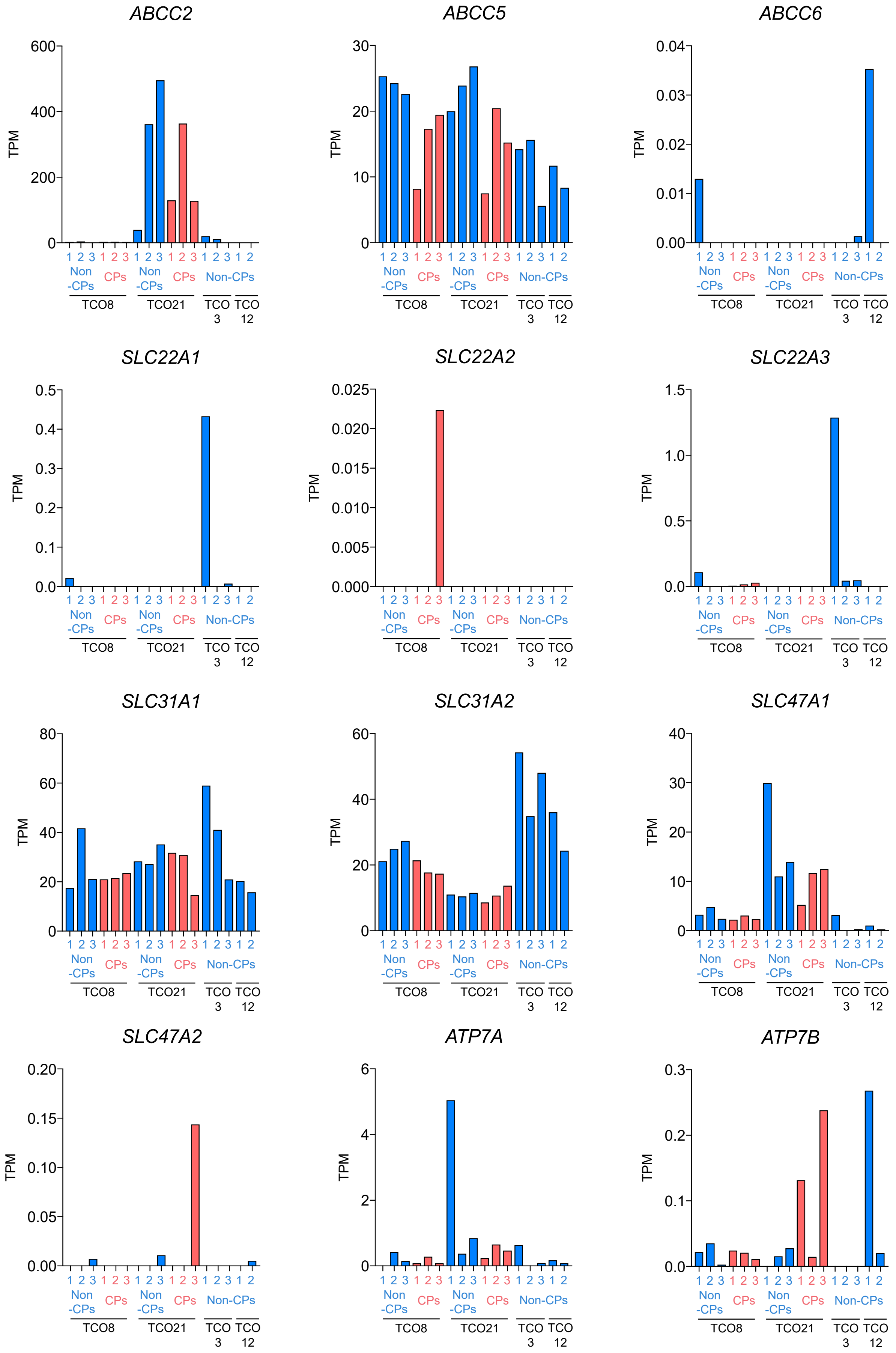

**Extended Data Fig. 2. Gene expression profiles of CDDP transporters in CPs and non-CPs (related to Fig. 5).** Gene expression of 12 transporters associated with either uptake or efflux of CDDP in CPs from TCO8 and TCO21 or in non-CPs from TCO8, TCO21, TCO3, and TCO12 are shown. Data are reported as TPM of RNA-seq data.

**Extended Data Fig. 3. Sato *et al.***

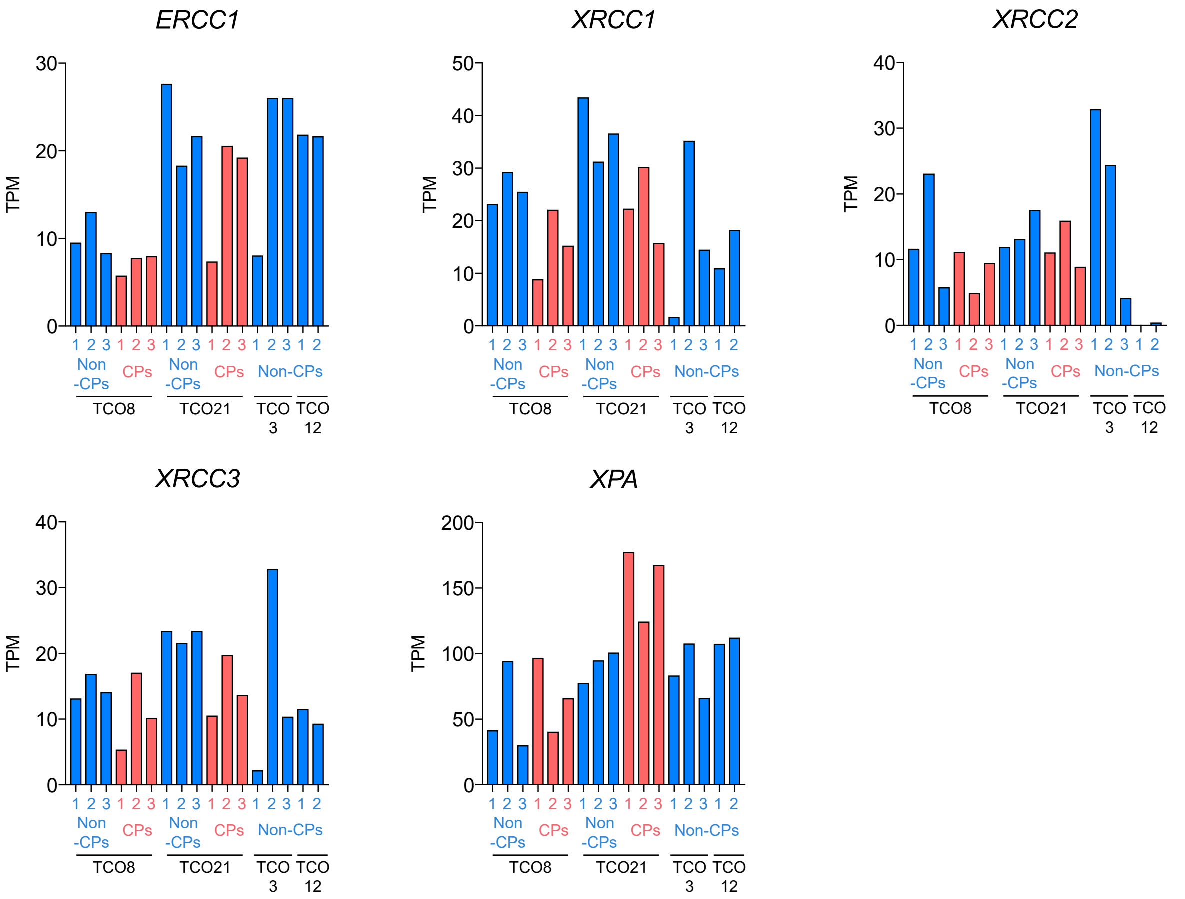


**Extended Data Fig. 3. Gene expression profiles of DNA repair enzymes for platinum drug adducts in CPs and non-CPs (related to Fig. 5).** Gene expression of 5 DNA repair enzymes for platinum drug adducts in CPs from TCO8 and TCO21 or in non-CPs from TCO8, TCO21, TCO3, and TCO12 are shown. Data are reported as TPM of RNA-seq data.

**Extended Data Fig. 4. Sato *et al.***

**a**

| sample | total_peaks | total_tags | tags_in_peaks | FRiP_percent |
|---|---|---|---|---|
| TCO8 rep.1 | 173285 | 88479167 | 34572006 | 39.07 |
| TCO8 rep.2 | 136223 | 49123655 | 18981133 | 38.64 |
| TCO21 rep.1 | 155300 | 65899039 | 28222575 | 42.83 |
| TCO21 rep.2 | 163686 | 62401345 | 27041648 | 43.34 |

**b**

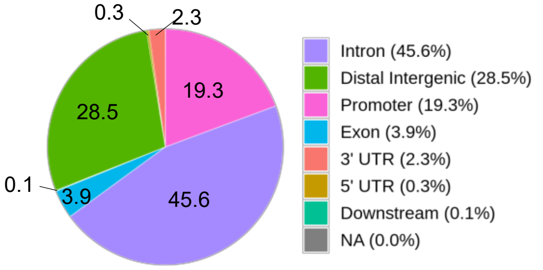


**c**

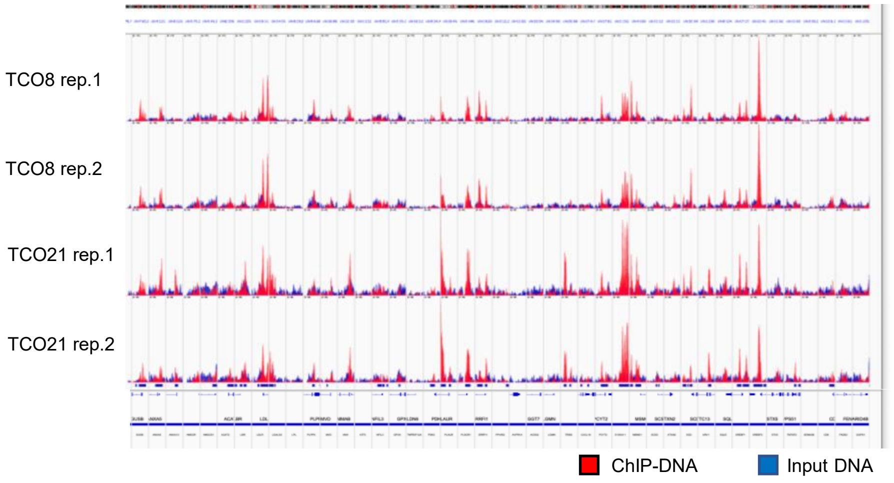


**d**

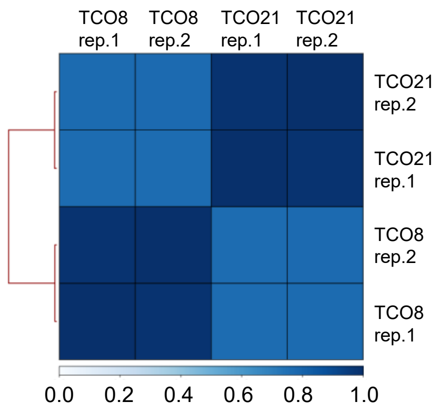


**e**

| Rank | Motif | P-value | log P-pvalue | % of Targets | % of Background | STD(Bg STD) | Best Match/Details | Motif File |
|---|---|---|---|---|---|---|---|---|
| 1 | 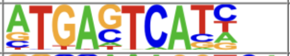 | 1e-14912 | -3.434e+04 | 25.06% | 3.95% | 48.6bp (61.0bp) | FOS::JUNB/MA1134.2/Jaspar(0.993) More Information \| Similar Motifs Found | motif file (matrix) |
| 2 | 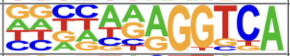 | 1e-2786 | -6.417e+03 | 23.58% | 11.97% | 49.7bp (58.5bp) | COUP-TFII(NR)/K562-NR2F1-ChIP-Seq(Encode)/Homer(0.976) More Information \| Similar Motifs Found | motif file (matrix) |
| 3 | 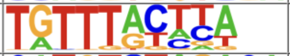 | 1e-2322 | -5.348e+03 | 16.66% | 7.73% | 51.6bp (57.7bp) | FOXM1(Forkhead)/MCF7-FOXM1-ChIP-Seq(GSE72977)/Homer(0.963) More Information \| Similar Motifs Found | motif file (matrix) |
| 4 | 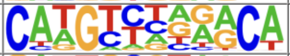 | 1e-1183 | -2.725e+03 | 9.64% | 4.62% | 50.6bp (61.1bp) | p53(p53)/Saos-p53-ChIP-Seq(GSE15780)/Homer(0.850) More Information \| Similar Motifs Found | motif file (matrix) |
| 5 | 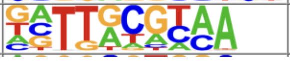 | 1e-993 | -2.287e+03 | 16.67% | 10.37% | 53.0bp (58.5bp) | NFIL3(bZIP)/HepG2-NFIL3-ChIP-Seq(Encode)/Homer(0.956) More Information \| Similar Motifs Found | motif file (matrix) |

**Extended Data Fig. 4. Summary and quality assessment of ChIP-seq data for transcription factor NR2F1 (related to Fig. 6f). (a)** Summary table showing ChIP-seq metrics for each sample, including the number of called peaks (total_peaks), total mapped reads (total_tags), number of reads within peaks (tags_in_peaks), and FRiP score (FRiP_percent), an indicator of signal enrichment. **(b)** Peak annotation distribution of NR2F1 binding sites, categorized by genomic regions. The pie chart displays the proportion of peaks located in each region. **(c)** Representative genome browser (IGV) snapshots showing typical ChIP-enriched regions. Tracks for NR2F1 ChIP (red) and input control (blue) are displayed for comparison, highlighting specific peak regions. **(d)** Heatmap of pairwise Pearson correlation coefficients (10-kb bin resolution) between biological replicates of TCO8 and TCO21, demonstrating reproducibility of the ChIP-seq experiments. **(e)** Top five de novo motifs identified by HOMER within $\pm$100 bp of peak centers. The table includes motif logos, statistical significance (p-values), and best matches to known motifs. Enrichment of the canonical binding motif of NR2F1 supports the specificity of the ChIP experiment.

### Extended Data Fig. 5. Sato *et al.*

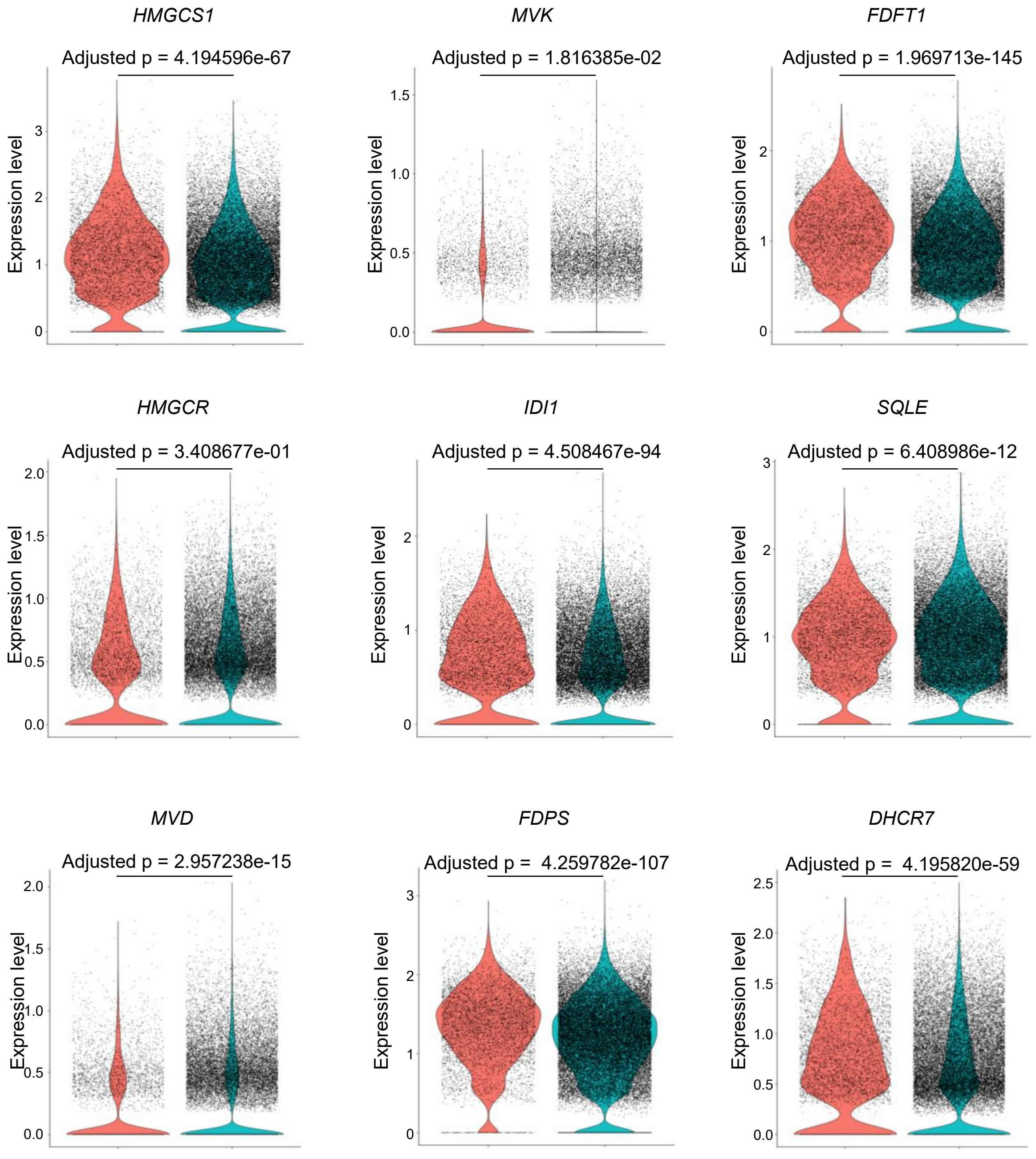

**Extended Data Fig. 5. NR2F1-mediated transcriptional regulation is associated with genes involved in the cholesterol biosynthetic pathway (related to Fig. 6g).** scMultiome analysis was performed on two chemo-resistant lines (TCO8, TCO21) and two chemo-sensitive lines (TCO3, TCO25), followed by integration of the snRNA-seq data. Violin plots showing expression levels of each cholesterol biosynthesis pathway-related gene in NR2F1-high and NR2F1-low cells defined in Fig.6d. Statistical analysis was performed using "MAST" method with bonferoni correction in the "FindMarkers" function in "Seurat" package.
